# Past viral infections can shape inter-individual variability in anti-viral TLR responses

**DOI:** 10.1101/2025.09.15.675924

**Authors:** Chad Hogan, Zafiirah Baurhoo, Nathália Beller, Ifeoma Osakwe, Ernest Turro, Samira Asgari

**Affiliations:** Institute for Genomic Health, Icahn School of Medicine at Mount Sinai, New York, NY; Department of Genetics and Genomic Sciences, Icahn School of Medicine at Mount Sinai, New York, NY; The Mindich Child Health and Development Institute, Icahn School of Medicine at Mount Sinai, New York, NY; The Charles Bronfman Institute for Personalized Medicine, Icahn School of Medicine at Mount Sinai, New York, NY

## Abstract

Viral infections can shape adaptive immunity, but whether they also influence innate immune responses later in life remains incompletely characterized. Here, we profiled serum from 12 healthy adults using VirScan, a high-throughput serological assay that infers prior viral exposures by mapping antiviral antibody reactivity across a broad range of viral epitopes. We paired this with *ex vivo* stimulation of TLR3 and TLR7/8 antiviral pathways and tested whether post-stimulation inflammatory responses are associated with prior viral exposures. At the antibody level, we observed consistent immunodominance at the protein level across individuals, alongside substantial variability at the epitope level. At the functional level, prior exposure to HSV-1, HSV-2, and norovirus was associated with differential production of CCL4, MCP-2, and TNF following TLR7/8 stimulation, respectively. This pilot study establishes a framework for integrating broad viral serology with functional immune profiling to investigate links between lifetime exposures and immune variability. Our results suggest that past viral infections, including acute infections, can contribute to variation in innate immune responses later in life. Larger studies will be required to validate these associations, establish causality, and determine the underlying mechanisms.

## Introduction

It is well established that past viral infections shape long-term adaptive immunity, primarily through neutralizing antibodies ^1^ and long-lived memory B and T cells ^2,3^. Less is known about how they shape long-term innate immune responses.

Current knowledge is largely restricted to a small number of studies on chronic infections, suggesting that viruses such as human immunodeficiency virus (HIV) and Epstein-Barr virus (EBV) can alter innate immune cells through lasting epigenetic, transcriptional, and metabolic changes that affect homeostasis and subsequent immune responses ^2,4,5^. More recently, acute infection with SARS-CoV-2 has been suggested to have similar lasting effects ^6^. However, existing evidence remains limited in both the range of viruses examined and the breadth of innate pathways studied, leaving the broader role of viral infections in shaping innate immunity through life incompletely understood. Approaches capable of detecting prior infection with many viruses, paired with broad profiling of innate immune responses, enable systematic identification of virus-innate immune associations and generation of hypotheses for mechanistic follow-up.

To this end, we conducted an exploratory pilot study (n = 12) combining high-throughput viral antibody profiling using VirScan ^7^ with proteomic profiling using Olink to assess whether prior viral infections are associated with inflammatory responses following *ex vivo* stimulation of Toll-like receptor (TLR) 3 and TLR7/8 antiviral pathways. We developed a rigorous analysis framework to reconstruct individual viral exposure histories, characterize antibody-based immunodominance, and link past viral exposures to TLR-mediated inflammatory responses, revealing virus-specific signatures for both chronic and acute infections.

## Results

### Serological reconstruction of viral infection history

To reconstruct individuals’ viral infection histories, we applied VirScan to serum samples from 12 individuals (**Table 1, Fig. S1**) ^7^. VirScan is a phage immunoprecipitation sequencing (PhIP-Seq) assay that detects antiviral antibodies to over 1,000 human-tropic viruses (**Table S1**), producing peptide-level antibody enrichment scores.

**Table 1:** Blood donors’ demographics and complete blood count (CBC) results.

| Donor | Age | Sex | Race*<sup>†</sup> | Ethnicity*<sup>†</sup> | WBC x10<sup>3</sup> per uL | RBC x10<sup>6</sup> per uL | Lymphocytes x10<sup>3</sup> per uL | Monocytes x10<sup>3</sup> per uL |
| --- | --- | --- | --- | --- | --- | --- | --- | --- |
| D1 | 43 | M | White | Non-Hispanic | 7.05 | 4.89 | 2.04 | 0.67 |
| D2 | 42 | F | White | Non-Hispanic | 6.86 | 5.11 | 2.14 | 0.46 |
| D3 | 63 | F | White | Non-Hispanic | 9.90 | 4.55 | 3.09 | 0.70 |
| D4 | 45 | F | Other | Puerto Rican | 8.50 | 4.30 | 2.19 | 0.55 |
| D5 | 35 | F | Other | Not reported | 4.81 | 4.13 | 2.42 | 0.30 |
| D6 | 58 | F | Black or African-American | Non-Hispanic | 6.99 | 5.31 | 2.78 | 0.48 |
| D7 | 67 | M | Other | Ecuadorian | 4.61 | 5.39 | 1.41 | 0.48 |
| D8 | 51 | F | White | Non-Hispanic | 5.64 | 4.32 | 1.49 | 0.31 |
| D9 | 67 | F | White | Not reported | 7.12 | 5.25 | 2.06 | 0.39 |
| D10 | 65 | M | White | Non-Hispanic | 9.49 | 3.24 | 2.63 | 0.56 |
| D11 | 62 | F | Bangladeshi | Non-Hispanic | 8.82 | 4.91 | 2.80 | 0.54 |
| D12 | 48 | F | White | Non-Hispanic | 4.78 | 4.93 | 1.68 | 0.50 |
| <sup>†</sup> Race and ethnicity were self-reported. |  |  |  |  |  |  |  |  |

While VirScan offers unmatched breadth and sensitivity, its outputs are susceptible to noise from non-specific binding and cross-reactive signals ^8^; robust interpretation therefore depends on careful data pre-processing, an aspect that has received uneven attention to date. We compared four peptide-to-virus mapping strategies and selected the one requiring two post-QC peptides mapping to the same viral protein from the same strain to call a "true" past infection. This minimized false positives while capturing known infections at expected epidemiological frequencies; less stringent thresholds inflated seropositivity to rare viruses (e.g., dengue), and stricter ones reduced detection of common viruses (e.g., RSV, **Fig. S2**).

Across individuals, we detected antibodies to an average of 290 viral peptides and 118 proteins spanning 26 human-tropic viruses (**Fig. 1A**), consistent with prior VirScan studies reporting 15-25 viruses per person ^7,9^. Common viruses such as EBV and Human Respiratory Syncytial Virus (RSV) were frequently detected, while less prevalent ones such as Hepatitis C Virus (HCV) appeared infrequently (**Fig. 1B, Table S3**). Prevalent viruses also yielded more enriched peptides per individual (**Fig. 1C, Table S3**).

**Fig. 1:**
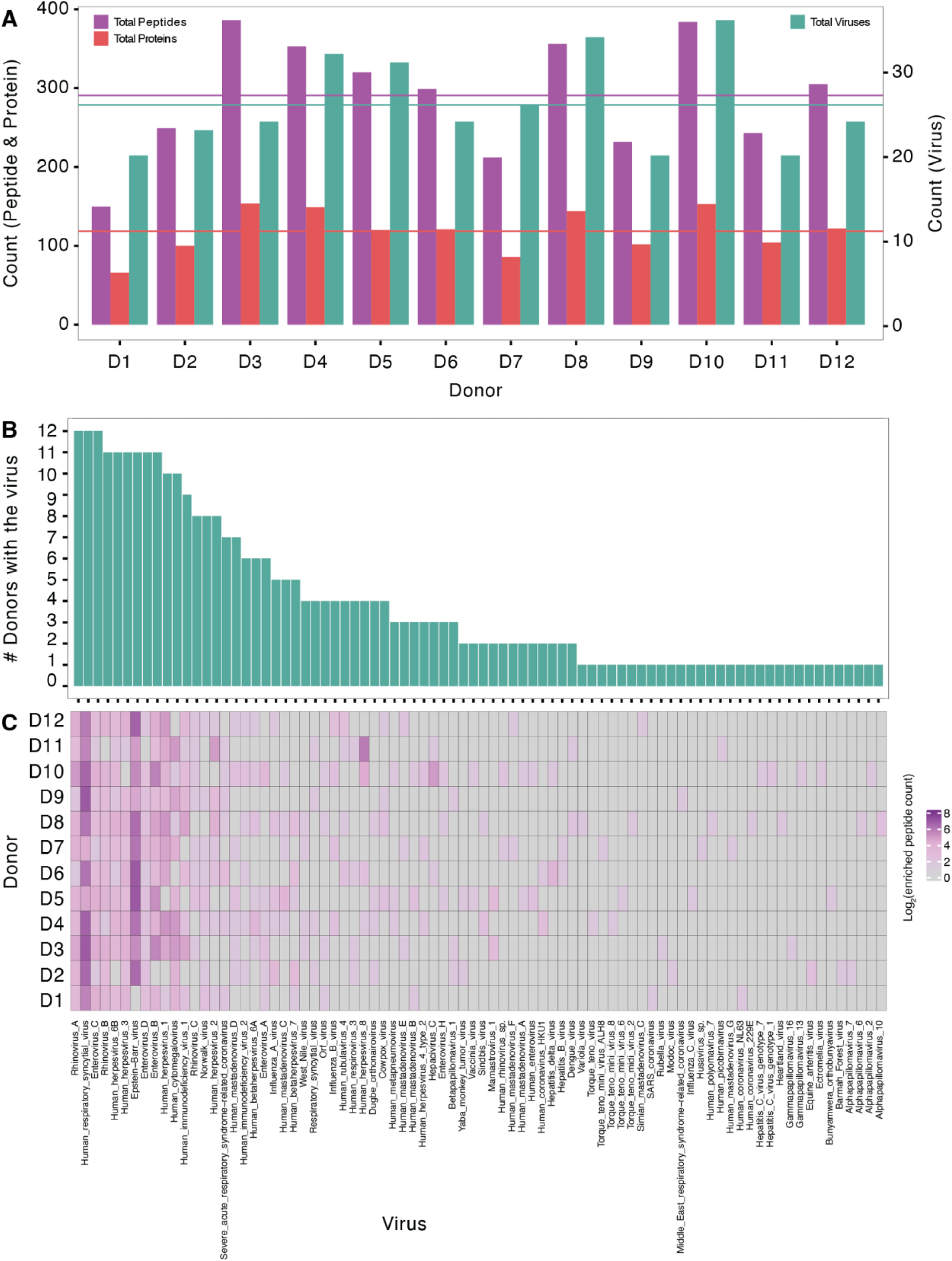
Profiling viral exposure histories in 12 healthy adults using VirScan. (**A**) Number of post-QC human-tropic viral peptides (purple), proteins (red), and viruses (turquoise) detected for each participant. A virus was considered present if ≥ 2 distinct epitopes from the same protein were enriched (log_2_FC > 2 and p < 0.0001). Horizontal lines indicate the cohort mean for each measure. (**B**) Virus prevalence across the cohort in descending order. (**C**) Number of viral peptides detected for each virus in each participant.

Despite careful QC, residual cross-reactivity may persist. For example, six individuals were HIV seropositive (**Fig. 1C**), yet their electronic health records showed no history of HIV. A plausible explanation is cross-reactivity with other viruses or human endogenous retroviral (HERV) elements, which share sequence similarity with HIV proteins ^10^. Such cross-reactivity is difficult to distinguish from true infection and may extend to other viruses with sequence homology; we therefore considered known epidemiology when interpreting downstream results.

### Antibody-based immunodominance and inter-individual variability

Immunogenicity varies between antigens and among epitopes within the same antigen, a phenomenon known as immunodominance, well studied for T cells but less so for antibody responses ^11^. Among viruses detected in five or more individuals, some proteins elicited responses in all donors, while others only in a subset. For RSV (present in 12 individuals), the attachment (G) glycoprotein elicited strong responses in every donor, followed by the fusion (F) protein (**Fig. 2A**), which are the only RSV proteins known to induce neutralizing antibodies ^12–14^. For influenza A virus (IAV; five individuals), all had antibodies to hemagglutinin (HA), followed by neuraminidase (NA) (**Fig. 2B**), consistent with established infection- and vaccine-induced responses ^15–17^.

**Fig. 2:**
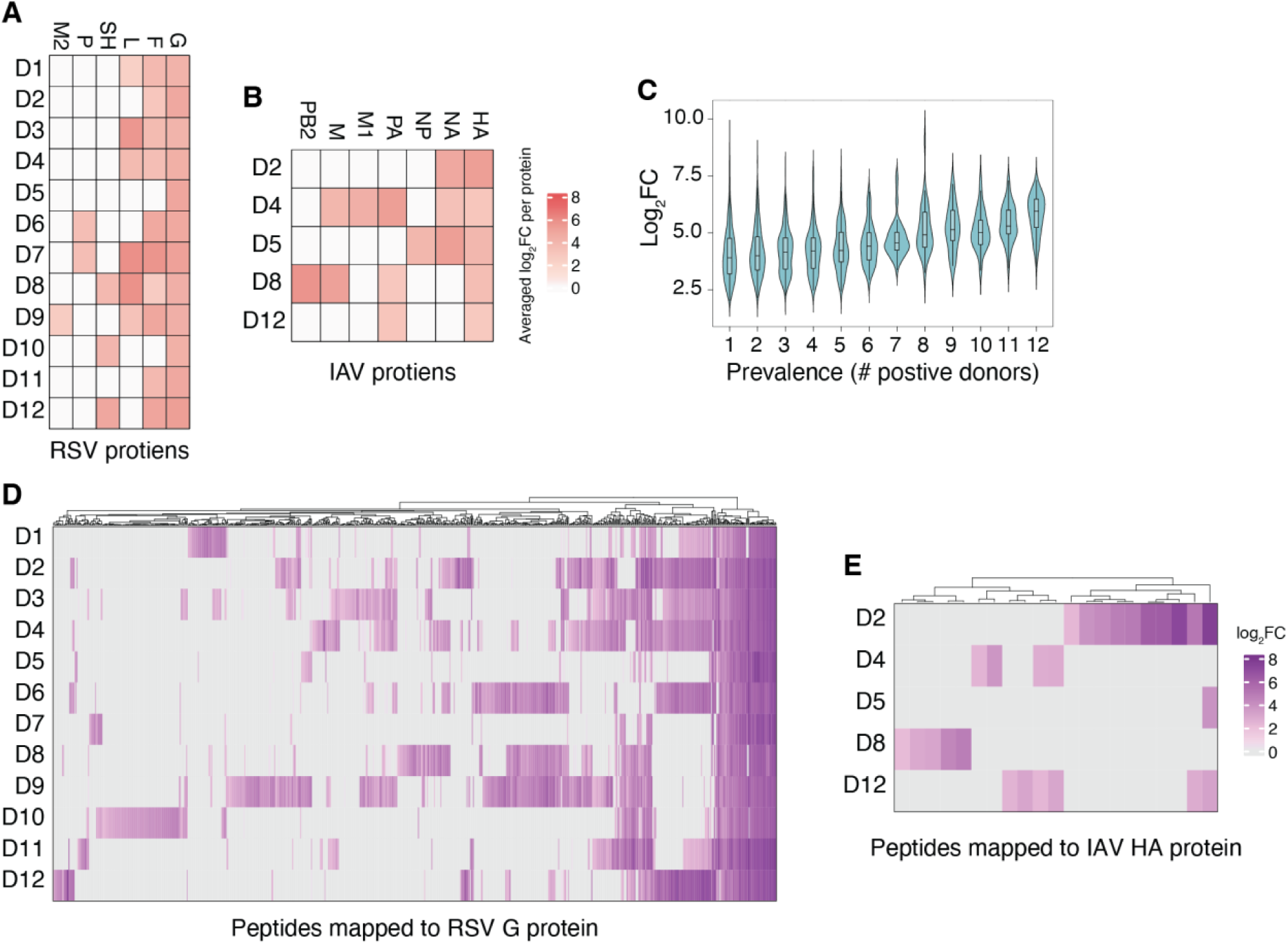
High-resolution antibody profiling against RSV and IAV. (**A & B**) Protein-level antibody reactivity results for RSV and IAV proteins. Each row represents a donor, each column a protein, colored according to the averaged log_2_FC of post-QC peptides mapped to that protein. Same proteins from the same virus with different UniProt IDs were grouped for visualization. ( **C**) Correlation between peptide enrichment values and their prevalence in our cohort (Pearson correlation coefficient (*r*) = 0.25, p-value < 2.2x10^−1^^6^) across all post-QC peptides mapped to at least one individual. More prevalent peptides were also more enriched compared to the background. (**D & E**) Peptide-level enrichment patterns for RSV attachment (G) proteins (812 post-QC peptides that were present in at least one donor) and IAV hemagglutinin (HA) proteins (29 post-QC peptides that were present in at least one donor). Each row represents a donor, each column a post-QC peptide, colored according to log_2_FC relative to beads-only decoy samples. These results reflect antibody-based immunodominance of certain viral proteins and peptides.

Peptide-level variability was greater. Peptides with stronger enrichment were detected in more donors and showed lower variability (**Fig. 2C**). Within proteins, recognition was highly heterogeneous: of 1,893 post-QC RSV G peptides, 812 (42.9%) were detected in at least one donor but only 60 (3.2%) in all donors (**Fig. 2D**). This likely reflects sequence diversity in antibody-targeted regions enabling RSV immune evasion ^18^; consistent with this, broadly recognized peptides mapped to the conserved central domain of G (**Fig. S3**) ^18^. A similar pattern was seen for IAV HA, where most peptides were recognized in only one individual (**Fig. 2E**), consistent with high mutation rates that facilitate immune escape ^19^. Comparable trends were observed for HSV-1, HSV-2, EBV, and Enterovirus B (**Fig. S4**).

### Cytokine responses to TLR stimulation and association with past viral exposures

To assess whether prior viral infections influence later antiviral innate responses, we stimulated whole blood from the 12 donors with TLR3 and TLR7/8 agonists, measuring 92 cytokines at 4 and 20 hours post-stimulation. Sixty-five were detected in ≥90% of samples (N = 72). Principal component analysis showed PC1 separating stimulated from unstimulated samples, PC2 separating TLR3 from TLR7/8 stimulation, and PC3-PC4 stratifying by time (**Fig. 3A-B, Fig. S5**).

**Figure 3:**
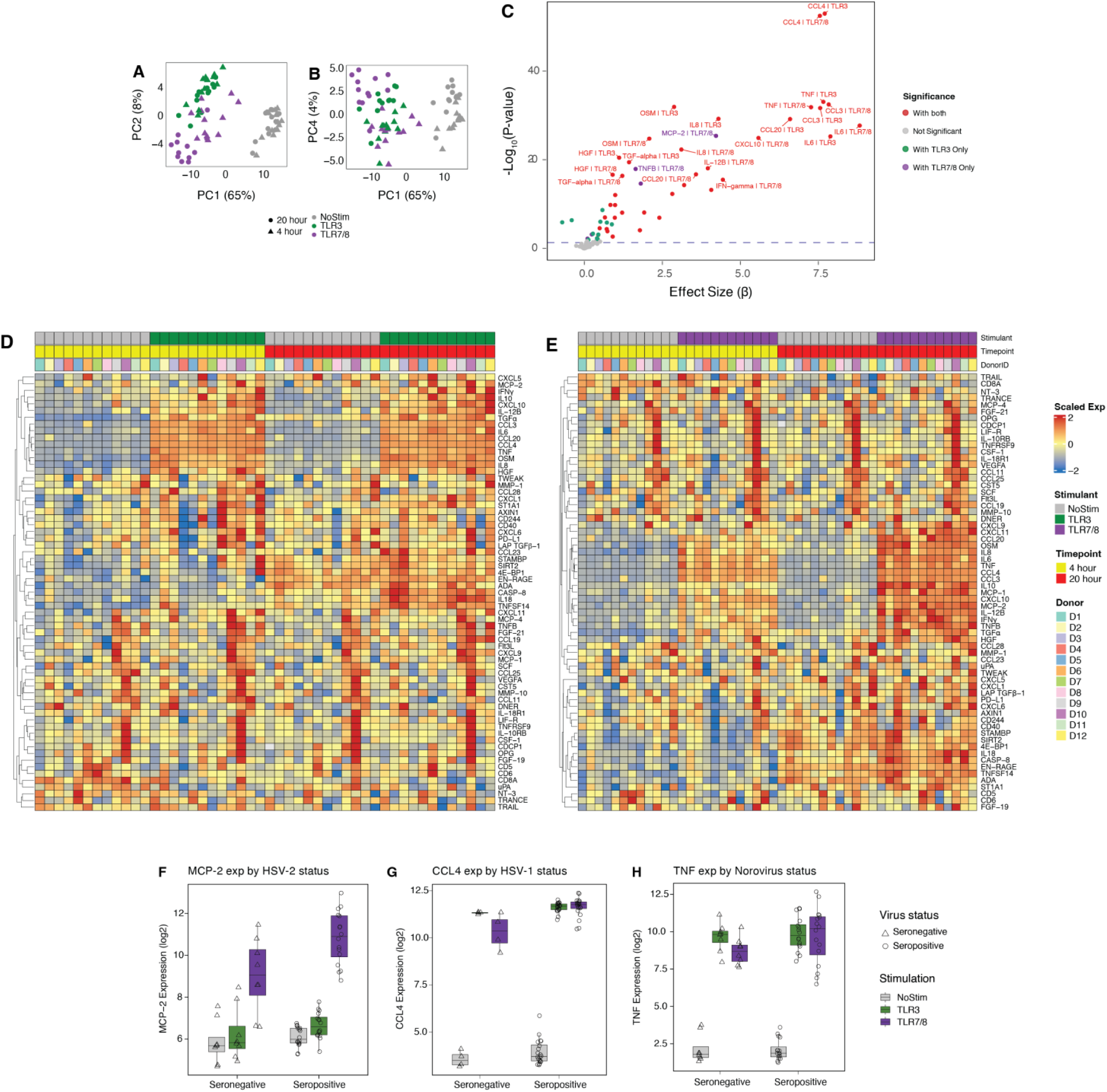
Cytokine responses reveal innate immune dynamics shaped by viral exposure. (**A & B**) Principal component analysis (PCA) of normalized protein expression (NPX) from unstimulated and TLR3- or TLR7/8-stimulated samples. PC1 and PC2 (panel A) separate non-stimulated samples from the stimulated ones. PC4 separate samples by time point (panel B). (**C**) Differential expression analysis results for normalized protein expression values under different stimulation conditions. X axis: effect size, Y axis: log_10_p-value. The horizontal dashed line represents FDR = 0.05. (**D & E**) Heatmaps of normalized expression values scaled across donors for cytokines under TLR3 and TLR7/8 stimulation compared with unstimulated controls. Normalized expression values were scaled across all individuals for each cytokine so that differences reflect relative rather than absolute levels. Colored bars on top of the heat maps represent stimulation condition, time point, and donor. (**F-H**) Significant associations (empirical p-value < 0.05) between the presence of specific viruses, ascertained using VirScan results, and normalized cytokine expression values following stimulation. Among which two pairs (HSV-1:CCL4 and HSV-2:MCP-2) showed significant interaction stimulation (FDR < 0.05).

Nineteen cytokines were differentially expressed under both stimulations (FDR < 0.05, 130 tests), 12 only after TLR3, and 6 only after TLR7/8 (**Fig. 3C, Table S4**). Canonical antiviral cytokines including CXCL10, IL-6, and TNF were upregulated, reflecting interferon-regulated pathways ^20,21^, alongside CXCL9 and IL-10, consistent with TLR-induced chemokine and regulatory responses ^21–23^ (**Fig. 3D-E**). Forty-one cytokines differed between 20 h and 4 h (FDR < 0.05, 65 tests; **Fig. S6, Table S5**), with most (33) increasing over time, consistent with myeloid activation ^24^ and the early inflammatory response within 24 h of infection ^25,26^. Nineteen cytokines showed stimulation-dependent changes with time (**Fig. S7, Table S6**).

To identify associations between past viral exposures and innate cytokine responses, we extended the differential expression model to include viral seropositivity status as a covariate, testing each virus-cytokine pair across stimulation conditions and time points. Of 2,176 tests, seven virus-cytokine pairs reached significance at FDR < 0.05 (**Table S7**). To guard against spurious associations driven by the small sample size and unbalanced seroprevalence, we further evaluated each pair using permutation testing (200 permutations per pair), which confirmed five associations at empirical p < 0.05 (**Table S8**). Of these, we excluded two based on large deviation of cohort seroprevalence from population seroprevalence: HIV-2, detected at far higher prevalence than expected (50% vs. <1%), and human metapneumovirus (hMPV), which is expected to be near-universal, was detected in only three individuals.

Among the remaining three pairs, HSV-1 was associated with elevated CCL4, HSV-2 with elevated MCP-2, and norovirus with elevated TNF (**Fig. 3F-H, Table S8**). HSV-1:CCL4 and HSV-2:MCP-2 also showed TLR7/8-specific interactions (**Fig. 3F-G, Table S9**). These associations remained consistent under stricter peptide-to-virus mapping, with detection supported by multiple peptides mapping to established viral proteins (**Fig. S7C-D, G-H**), increasing confidence that they reflect biologically meaningful prior exposures rather than cross-reactive signals. No significant interaction with time was detected (**Table S10**).

## Discussion

Traditionally, innate changes following viral infection have been viewed as transient, with long-term protection maintained by adaptive memory ^2^. This view is challenged by trained immunity, in which innate cells acquire memory-like features through durable epigenetic, transcriptional, and metabolic changes ^27^, documented for live attenuated vaccines such as BCG and MMR ^28,29^ and emerging for acute viral infections including influenza ^30,31^ and SARS-CoV-2 ^6^. While our cross-sectional design cannot directly demonstrate trained immunity, the virus-specific cytokine signatures we observe (HSV-1, HSV-2, and norovirus seropositivity each linked to distinct cytokine responses) are consistent with the possibility that prior infections leave durable imprints on innate immune function. Notably, we observed such signatures for both chronic and acute viruses, in line with emerging evidence that trained immunity is not restricted to chronic exposures.

Our findings also have implications for antibody-based immunodominance across the virome. Consistent recognition of the conserved central domain of RSV G aligns with reports identifying this region as a target for monoclonal antibodies and recombinant vaccines ^18,32^. Extending VirScan to larger cohorts will enable an unbiased study of immunodominance for chronic viruses such as HSV-1/2, EBV, and CMV, and the discovery of vaccine targets for common acute infections such as enteroviruses, rhinoviruses, and coronaviruses.

A key technical insight is that VirScan interpretation is highly sensitive to data processing, with filtering criteria at the epitope and protein levels strongly influencing prevalence estimates. Our findings underscore the need for standardized, context-specific filtering strategies to ensure reproducibility across studies ^33,34^.

Our study has several limitations. Given the cross-sectional design and limited sample size, observed differences may reflect baseline immune set points, latent or intermittent viral reactivation (particularly for herpesviruses), or residual confounding. The *ex vivo* system captures only a subset of pathways activated during natural infection ^35–38^, and VirScan has known constraints, including antibody waning, reliance on linear peptides, and cross-reactivity ^33^. Other determinants of immune variation, including host genetics, microbiome, and environment, will require substantially larger cohorts to integrate meaningfully.

In summary, our study presents an integrated, rigorous framework for systematically dissecting how lifetime viral exposures shape human immunity. Applying such approaches at scale is increasingly important given growing epidemic and pandemic threats, and may ultimately inform personalized vaccines, antivirals, and pandemic preparedness.

## Methods

### Study participants

We recruited donors from the Mt. Sinai Hospital system. All were adults between the ages of 18 and 70. Donors were generally ‘healthy’, had no aspirin or antiplatelet drug intake in the last five days, no steroid or anti-inflammatory drug intake in the last 24 hours, no fever in the past 24 hours, no myocardial infarction or stroke in the past year, no blood, platelet, or coagulation factor transfusions in the last month, no history of bone marrow transplantation, and a body mass index (BMI) below 35. We obtained informed consent from all donors for blood collection and consequent analysis following a protocol that was approved by Mount Sinai’s Internal Review Board (STUDY ID# 22-01173). In total, we processed blood from 12 individuals for this study. Briefly, we collected up to 4 mL of blood into EDTA-treated tubes. From these tubes, we used ∼50 μL to perform complete blood counts (CBCs) using a Sysmex XN-10 hematology analyzer within 2 hours of the time of collection. We used the rest for VirScan analysis and immunostimulation as described below.

### VirScan analysis using Phage Display Immunoprecipitation Sequencing (PhIP-Seq)

For each donor, we collected serum from 250 μl of whole blood processed using the standard centrifuge protocol to perform VirScan. VirScan is a high-throughput serological profiling platform based on programmable phage immunoprecipitation sequencing (PhIP-Seq) that enables broad, epitope-level characterization of antiviral antibody repertoires ^7^. The method uses a synthetic library of bacteriophages displaying short, overlapping peptides that collectively span the proteomes of nearly all known vertebrate viruses. When incubated with human plasma or serum, antibodies bind to cognate peptide epitopes, and antibody-bound phages are subsequently immunoprecipitated and identified by next-generation sequencing. The resulting read counts provide a quantitative measure of antibody reactivity against individual linear viral epitopes.

VirScan services were provided by CDI Labs Canada (Toronto, Canada). The VirScan library comprises over 480,000 62-mer overlapping peptides from nearly all known vertebrate viruses with 14-mer amino acid adjacent overlaps as described previously ^7^. Library cloning, sample screening, PCR, and peptide read count data curation were performed as described in Mohan, D., et al. ^39^

Briefly, screens were performed in 0.5 mL PBS containing a phage library at an average of 10^5^ excess clones per unique phage in the input library (approx. 4.8 × 10^10^ total input plaque-forming units) with 0.2 μL serum (approx. 2 μg IgG). The mixture was rotated overnight to allow antibodies to bind any phage-displayed peptide targets. The next day, a Protein A/G coated magnetic bead slurry was added to some samples and rotated for an additional 4 hours to capture all IgG. All protein A/G beads were then washed and resuspended with a Herculase II Fusion Polymerase master mix and amplified with PCR. Protein A/G is a recombinant fusion protein that combines the antibody-binding domains of both Protein A and Protein G. This combination results in a molecule with a broad binding affinity for various immunoglobulin (IgG) subclasses. These ‘beads-only’, mock-immunoprecipitation buffer-alone samples were included to control for non-specific binding. Positive controls included samples that were spiked with a polyclonal antibody against glial fibrillary acidic protein (GFAP) human protein and its isoforms. Anti-GFAP antibodies are not typically found in human plasma or serum samples. GFAP peptides are included in the PhIP-seq library and thus are expected to be found in the spiked-in samples at high levels. Following the wash, multiplex barcodes and sequencing adapters were then incorporated during a subsequent PCR reaction. Barcoded samples were pooled and sequenced using Illumina instruments to obtain paired-end reads of the amplified phage inserts. Paired-end reads were demultiplexed, trimmed, and aligned to the viral peptide library using bwa mem, requiring a minimum alignment score of 50 (“-T 50” argument). Reads aligning to each phage clone sequence were counted and recorded.

To assess antibody reactivity enrichment, we performed differential expression analysis using the edgeR exactTest function to estimate fold changes (FC) and p-values for each sample against the ‘beadsonly’ decoy samples. We considered epitopes significantly enriched if they had a log_2_FC of 2 or higher over the ‘beadsonly’ controls (i.e., at least 4 times enriched) and an unadjusted p-value of 10^−4^ or lower. This p-value threshold was chosen based on CDI Labs’ recommendations to minimize false positives. For comparison, this is a more stringent threshold than an FDR-adjusted p-value < 0.05 for the total number of tests performed (i.e., p-value < 0.0011 for N = 1,242,258 tests). Finally, we removed peptides derived from non-human-tropic viruses according to CDI Lab’s labeling of the possible hosts for each virus. This step improves robustness and biological interpretability. Given the small size of our study, non-human viruses that pass all filtering criteria are likely to reflect antibody cross-reactivity, environmental exposure, or microbiome-derived signals rather than true human infection. We kept all significantly-enriched peptides that were derived from viruses that can infect humans, removing only those that did not have humans listed as a potential host, such as bacteriophages and baculoviruses or viruses known to infect non-human vertebrates only. We refer to enriched peptides that pass these filtering criteria as “post-QC peptides”.

### Mapping of viral peptides to viral proteins

To classify peptides according to their source proteins, we extracted the UniProt accession identifiers included in the VirScan library annotations for each peptide (provided by CDI labs). We then queried these accession numbers against the UniProt database to obtain the corresponding protein information, including virus name, protein name, and viral strain. To calculate per-protein log_2_FC values, we averaged log_2_FC values of all the peptides that were mapped to a protein that had the same name in the same virus, regardless of viral strain.

### Mapping of viral peptides to viruses

To ascertain whether a virus is detected in a sample or not, we mapped post-QC peptides to viruses. This mapping can be done at different stringency levels to determine what is a confidently detected virus. To determine the optimal approach for collapsing peptide-level data into virus-level calls, we systematically evaluated four strategies:

Option 1: A virus was considered present in a sample if at least one peptide from any viral protein from any viral strain was enriched (i.e., log_2_FC > 2 and p-value < 10^−4^ in the edgeR exactTest function).
Option 2: Similar to option 1 but requiring that at least two peptides per virus were enriched, regardless of whether they came from the same or different viral proteins and strains.
Option 3: A virus was called present if at least two enriched peptides were mapped to the same viral protein and the same viral strain (i.e., mapped to the same viral protein UniProt ID).
Option 4: Similar to option 3 but requiring three or more enriched peptides mapped to the same viral protein and strain.

These strategies were evaluated to balance sensitivity (detecting true positives) and specificity (minimizing noise or spurious calls). This analysis demonstrates that virus detection is highly sensitive to the peptide filtering threshold and highlights the importance of carefully selecting a consistent and biologically interpretable rule for defining viral hits.

We selected Option 3, which requires at least two enriched peptides from the same viral protein and the same strain for all downstream analyses. This approach offered a practical balance of robustness and interpretability, while also aligning well with known epidemiological infection prevalences and donors’ electronic health records (EHRs) available.

### Mapping peptides to the RSV G protein

Post-QC peptides corresponding to the Glycoprotein G (G protein) of Human Respiratory Syncytial Virus (RSV), which were detected in all 12 donors, were selected for positional mapping. The reference full-length RSV G protein sequence (strain A2, UniProt accession P03423, 298 amino acids) was retrieved from the UniProt database. The selected peptide sequences were compiled into a FASTA file and aligned to the full-length RSV G protein using BLASTp (NCBI web interface) with default settings.

### Immune stimulation

Venous blood was used within 4 h of collection. The assay was performed in 2-ml polypropylene tubes, to which 500 μl of whole blood was added. The stimulation assay was performed by adding to all tubes 1000 μl RPMI-1640 medium (Gibco, cat: 11875093), penicillin (100 U/mL), streptomycin (100 μg/mL) L-glutamine (2 mM)( Gibco, cat: 10378016). The tubes were incubated either without stimulus or in the presence of Toll-Like Receptor (TLR)-7/8 (Resiquimod, R848, conc: 0.3 μg/mL; Sigma-Aldrich, St. Louis, MO, USA; catalog no. SML0196) or TLR-3 (Polyinosinic-polycytidylic acid, Poly(I:C), conc: 20 μg/mL; Sigma-Aldrich, St. Louis, MO, USA; catalog no. P1530) agonist at 37°C for the indicated time point. Stimulated samples were harvested at 4 hours and 20 hours post stimulation for the no stimulus condition as well as the TLR-3 and TLR-7/8 stimulated conditions, spinning down the sample at 1000 x G for 10 minutes and collecting the supernatant for proteomics analysis.

### Olink proteomics

We submitted the collected supernatant samples to the Human Immune Monitoring Core at the Icahn School of Medicine at Mount Sinai for protein quantification using Olink. Protein quantification was performed using the Olink® Target 96 Inflammation panel (Olink Proteomics, Uppsala, Sweden), based on Proximity Extension Assay (PEA) technology. The platform measures 92 inflammation-related proteins simultaneously, and results are reported as Normalized Protein eXpression (NPX) values on a log_2_ scale.

### Olink data Quality Control

Data preprocessing and initial quality control were performed by the Olink core facility, following standard protocols. Briefly, protein expression values were reported as Normalized Protein eXpression (NPX) values on a log_2_ scale, normalized against internal and inter-plate controls to account for technical variation. We reviewed the provided QC flags and confirmed that all samples passed internal quality thresholds. No samples were flagged for low detection rate or high variability. For each protein, we examined detection rates across the cohort and excluded proteins not detected in at least 90% of samples (N = 72 samples, corresponding to 12 samples, 3 conditions, and 2 time points). To further evaluate data quality, we visualized the distribution of NPX values across proteins and individuals and assessed inter-individual variance.

### Principal Component Analysis (PCA)

We used PCA for two purposes. First, we applied PCA to cytokine expression data to visualize inter-individual differences in response patterns. NPX values were log_2_-transformed by design and were mean-centered using the default setting in the prcomp function in R. We used these PCA analyses to visualize the broad and combined effects of stimulation and time point on protein expression across samples.

Additionally, we performed PCA on cytokine expression values from the unstimulated 4-hour samples to capture underlying differences in innate immune status across individuals prior to stimulation. These samples reflect each donor’s baseline cytokine profile. Similarly, we performed PCA on Complete blood counts (CBC) to account for variability in baseline blood composition. The following CBC components were included in the PCA: white blood cell count, red blood cell count, hemoglobin levels, platelet count, and absolute counts for the following immune cells: neutrophil, lymphocyte, monocyte, eosinophil, basophil, and immature granulocyte. Top PCs from these two analyses were included as covariates in downstream association models as a proxy for donors’ baseline immune status and to adjust for differences in the baseline immune status of donors.

### Differential protein expression analysis

We used a linear mixed model implemented in the lmer function in R to assess cytokine response to stimulation and time point as follows:

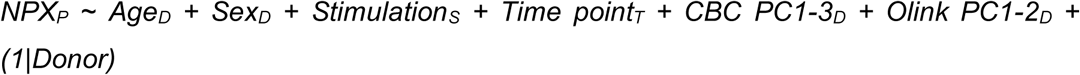

Where NPX is the normalized log_2_ scale protein expression value for protein P. Age and Sex, are donor-level fixed effects to capture differences in donors’ demographics. We also included 3 CBC PCs and 2 baseline Olink PCs as donor-level fixed effects to adjust for inter-individual immune variation unrelated to stimulation or viral history. The number of PCs was selected so they can capture > 60% of the observed variance in each modality. Stimulation is a fixed effect variable for stimulation condition S (categorical with three levels: no stim, TLR-3 agonist, TLR-7/8 agonist), and Time point is a fixed effect variable for time point condition T (categorical with two levels: 4h and 20h), 1|Donor is a random effect for donor.

Parameter estimation was performed using restricted maximum likelihood (REML). For stimulation analyses, “no stim” served as the reference group. Statistical significance was determined using the Benjamini-Hochberg (BH) procedure to control the false discovery rate (FDR). A cytokine was considered significantly differentially expressed under a given stimulation if the stimulation term had an FDR < 0.05. Likewise, for time point analyses, 4 h served as the reference group, and a cytokine was considered significantly differentially expressed over time if the time point estimate had an FDR < 0.05. FDR control provides a balance between limiting false positives and maintaining power for our study, given the small sample size, the large number of tests across three stimulation conditions and all measured cytokines, and the high correlation among cytokines due to shared regulatory pathways. More conservative methods, such as Bonferroni correction, would be overly stringent under these conditions.

To assess whether the effect of time on cytokine expression depended on stimulation, we extended the model to include a Stimulation × Time point interaction term:

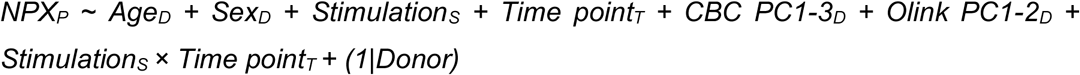

This interaction term tests whether the impact of stimulation on cytokine levels differs between the 4 h and 20 h time points (*i.e.* whether the effect of stimulation is dependent on time). All other model components were defined as above. Cytokines with a significant interaction term (FDR < 0.05) were interpreted as having time-dependent stimulation effects.

### Virus-protein association analyses

To test whether prior exposure to specific viruses affects the overall cytokine levels, we used a linear mixed model implemented in the lmer function in R with REML to estimate the parameters as follows:

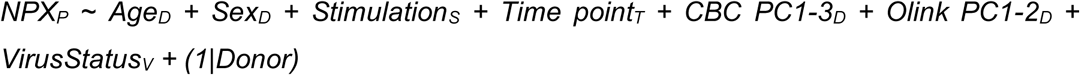

NPX, Age, Sex, Stimulation, Time point, CBC PCs, Olink PCs, and Donor are the same as described in the previous section. VirusStatus is a fixed effect variable for the presence or absence of virus V in each donor (categorical with two levels: detected and not detected). We restricted this analysis to viruses that were found in at least three individuals to avoid test statistic bias due to a small number of positive observations^40^. We then ran this model separately for each virus and used the BH procedure to determine statistical significance. Viruses with an FDR < 0.05 were considered significantly associated with cytokine levels.

To assess whether the effect of stimulation on cytokine expression depended on prior viral exposure, we extended the above mixed model to include a VirusStatus × Stimulation interaction term:

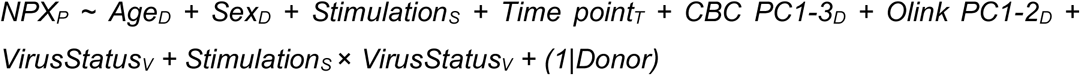

All other model components were defined as described before. Cytokines with significant interaction terms (FDR < 0.05) were interpreted as exhibiting differential responses to stimulation depending on whether the virus was detected in a donor.

Similarly, to determine whether the change in cytokine expression over time differed by viral status, we further extended the model to include a VirusStatus × Time point interaction term:

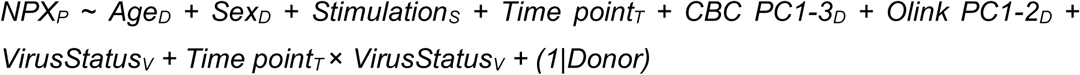

A significant interaction (FDR < 0.05) was interpreted as cytokine levels evolved differently over time in virus-positive versus virus-negative donors.

### Permutation analysis for virus-cytokine associations

To evaluate the robustness of virus-cytokine associations and ensure that significant findings were not driven by data structure or small numbers of seropositive individuals, we performed permutation analyses for each virus with at least one cytokine meeting the FDR < 0.05 threshold. For each virus-cytokine pair, we randomly reassigned virus status across individuals while keeping all other variables (stimulation condition, timepoint, demographic covariates, CBC PCs, Olink PCs), as well as the cytokine expression values, fixed. The total number of seropositive and seronegative individuals was preserved in every permutation.

For each virus, we generated 200 permuted datasets and refit the linear mixed model to obtain a distribution of association p-values under the null hypothesis of no relationship between virus status and cytokine expression. We used the empirical cumulative distribution function (ecdf in R) to compute an empirical p-value for each association by determining the percentile rank of the observed p-value relative to the permutation-derived null distribution. Virus-cytokine associations with empirical p-values < 0.05 were considered robust.

## Acknowledgements

This work was supported by a pilot award from Mount Sinai’s Institute for Genomic Health. Chad Hogan was supported by National Institute of Health, National Cancer Institute training award T32CA078207. Olink analyses were performed at the Human Immune Monitoring Center (HIMC) at Mount Sinai. We thank CDI Labs for providing technical expertise that aided in the analysis of PhIP-seq output and VirScan data. We are grateful to all study participants for their contribution to this work.

