## Supplementary figures for "Past viral infections can shape inter-individual variability in anti-viral TLR responses"

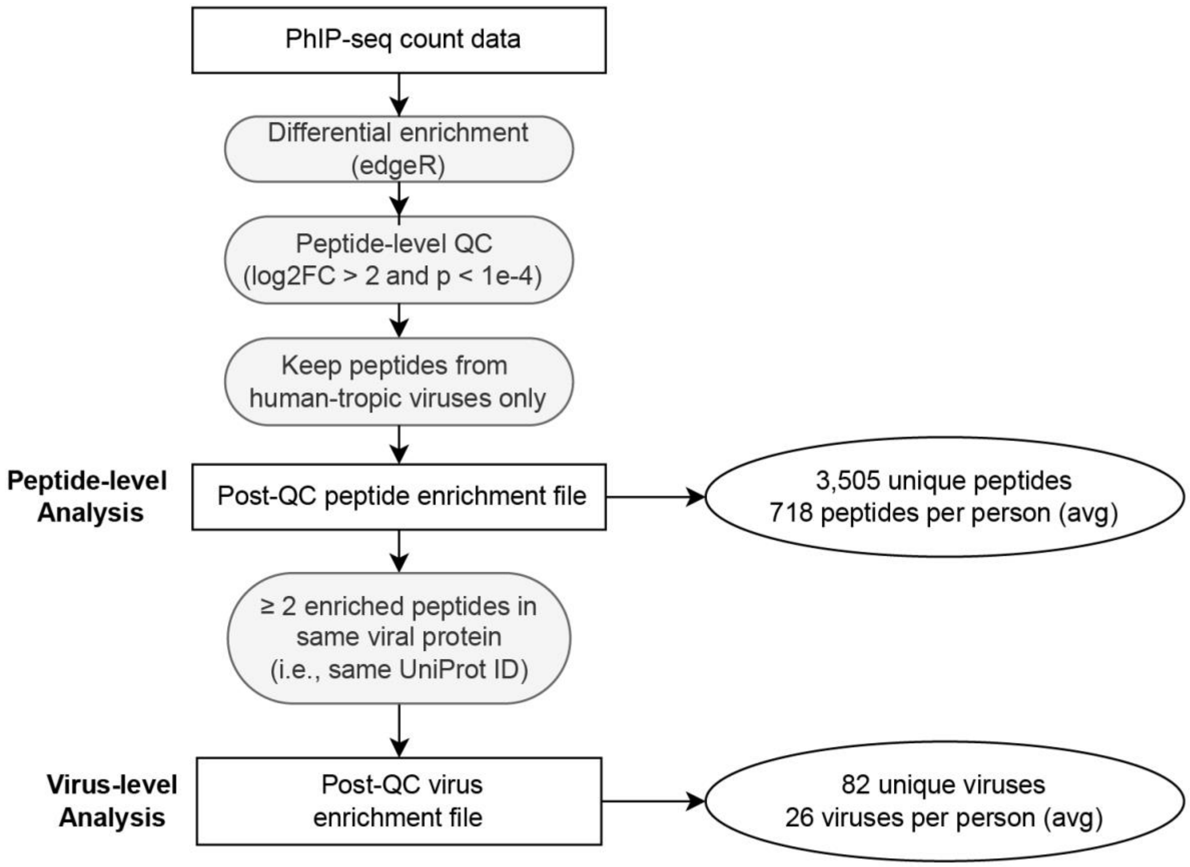


**Fig. S1: VirScan data analysis pipeline.** We used the above steps to identify viral peptides that were significantly enriched in samples, performed additional filtering to get to a high-confidence set of enriched peptides with high biological plausibility, and mapped those to viruses to determine whether a virus was detected or not in a donor.

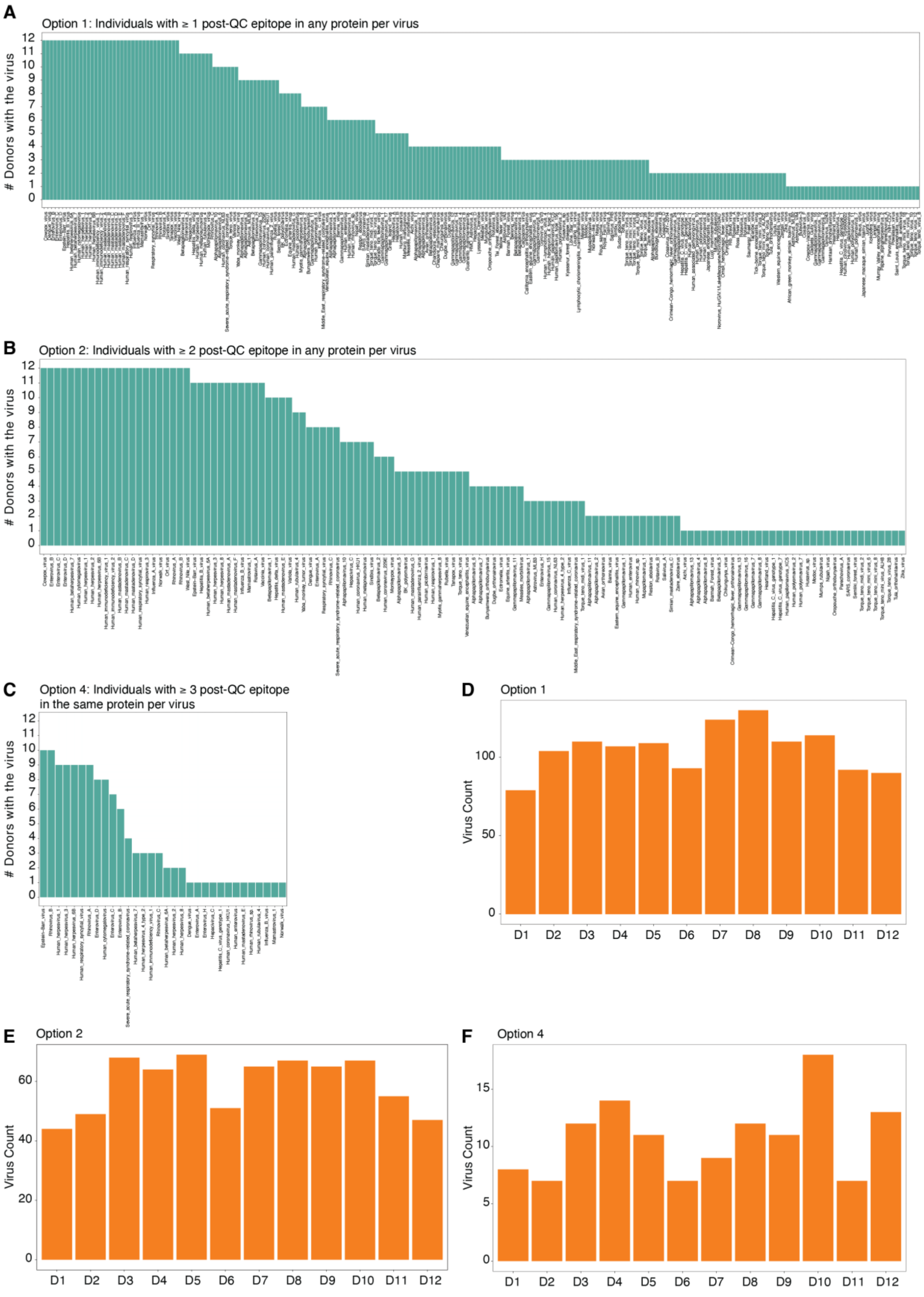
A
Option 1: Individuals with ≥ 1 post-QC epitope in any protein per virus
### Donors with the virus
B
Option 2: Individuals with ≥ 2 post-QC epitope in any protein per virus
### Donors with the virus
C
Option 4: Individuals with ≥ 3 post-QC epitope in the same protein per virus
### Donors with the virus
D
Option 1
Virus Count
D1 D2 D3 D4 D5 D6 D7 D8 D9 D10 D11 D12
E
Option 2
Virus Count
D1 D2 D3 D4 D5 D6 D7 D8 D9 D10 D11 D12
F
Option 4
Virus Count
D1 D2 D3 D4 D5 D6 D7 D8 D9 D10 D11 D12

**Fig. S2: Viruses found using different peptide-to-virus mapping strategies.** (**A & D**) Identified viruses and per donor virus counts using the least stringent option (Option 1), which considered a virus present in a sample if at least one post-QC peptide from any viral protein was present. This approach led to the detection of viruses such as cowpox and dengue virus (panel A), which are not expected to be prevalent in the population in all individuals, indicating a lack of sufficient control for false positives (possibly due to cross-reactivity of antiviral antibodies). In line with this, the total number of viruses found per donor (panel D) is 2-3 times higher than what is expected based on previous VirScan studies. (**B & E**) Similar to A and D for Option 2, which considered a virus present in a sample if at least two post-QC peptides from any viral protein were present. Although a smaller total number of viruses was found per donor (panel E), uncommon viruses such as cox pox and dengue virus (panel B) would be considered as present using this approach. (**C & F**) Similar to A and D for Option 4, the most stringent strategy. This approach considered a virus present in a sample if at least three post-QC peptides from the same viral protein were present in an individual. This approach underestimated the presence of antibodies against common recurring or chronic viruses that are expected to infect all or the majority of individuals, such as EBV, RSV, and Rhinovirus (panel C). Consequently, the total number of viruses detected per donor was also considerably lower than the 15-25 per person reported in previous VirScan studies (panel F). Based on these analyses and also comparisons with donors' medical records, we selected a strategy that required at least two post-QC peptides from the same viral protein to consider a virus present (option 3) for follow-up analyses (Fig. 1 in the main text).

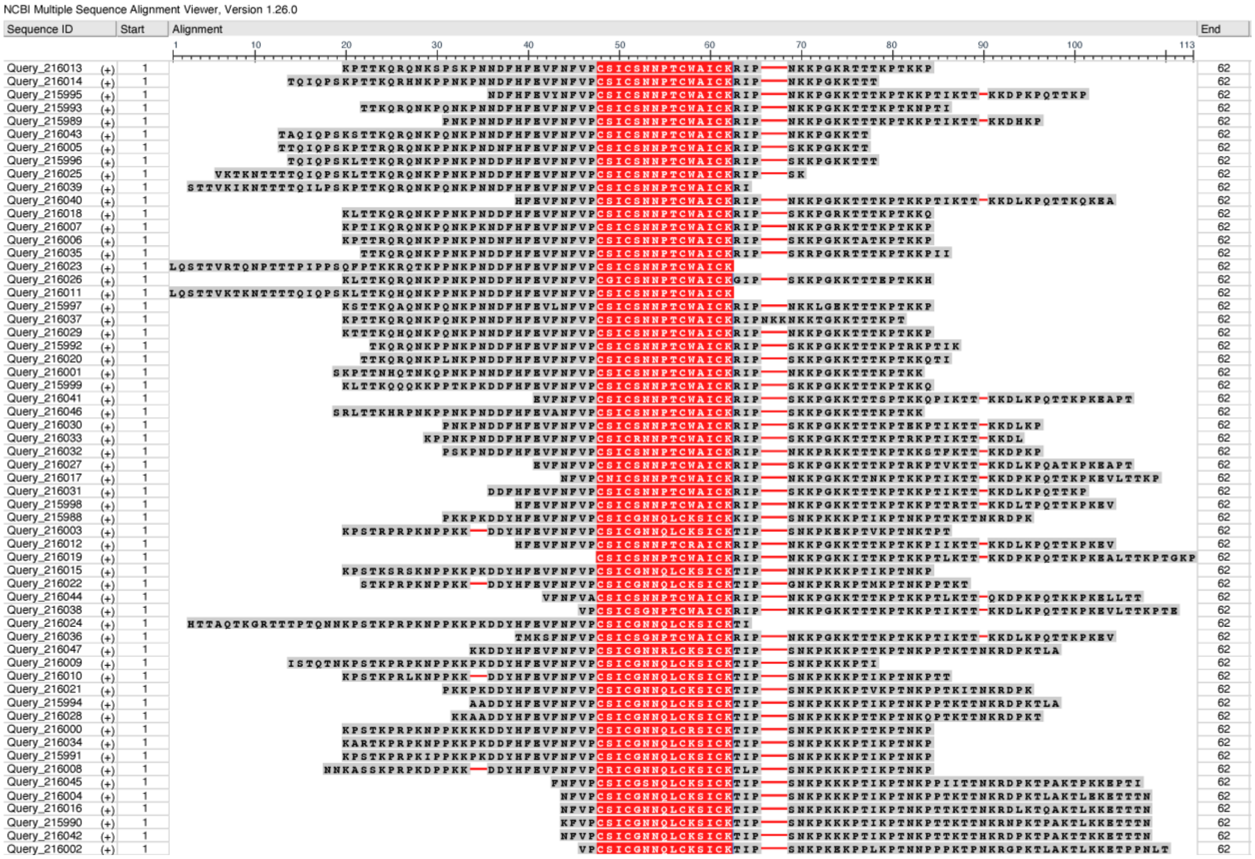


**Fig. S3. Alignment of RSV G protein peptides detected in all 12 participants.** Peptides map within the RSV G glycoprotein and cluster in the central conserved region (CCR). The amino acid sequence for the peptides is shown, with the red epitope sequence being the same across all peptides. These sequences match previously published G protein sequences from various strains (e.g. DOI: 10.1128/jvi.02201-21)

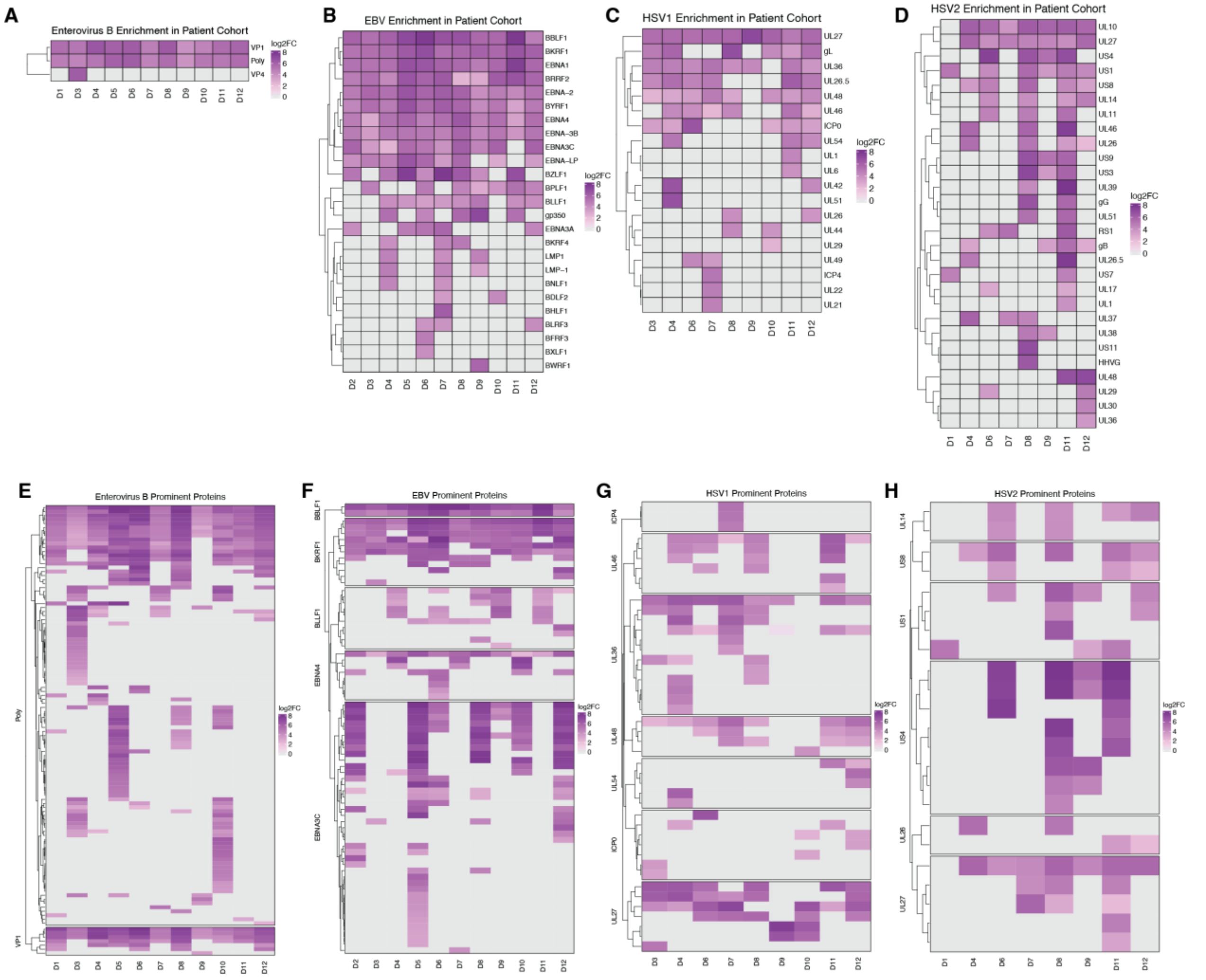


**Fig. S4. High-resolution antibody profiling for additional viruses.** (**A-D**) Protein-level enrichment ($\log_2$FC) for viral proteins from (**A**) Enterovirus B; (**B**) Epstein-Barr virus (EBV), (**C**) herpes simplex virus 1 (HSV-1), and (**D**) herpes simplex virus 2 (HSV-2). Each column represents a donor, each row a protein, colored according to the averaged $\log_2$FC of post-QC peptides mapped to that protein. Same proteins from the same virus with different UniProt IDs were grouped for visualization. (**E-H**) Peptide-level enrichment heatmaps for prominent proteins from each virus. (**E**) Enterovirus B capsid proteins. (**F**) EBV nuclear and structural antigens. (**G**) HSV-1 glycoproteins. (**H**) HSV-2 glycoproteins. Each column represents a donor, each row a post-QC peptide, colored according to $\log_2$FC relative to beads-only decoy samples.

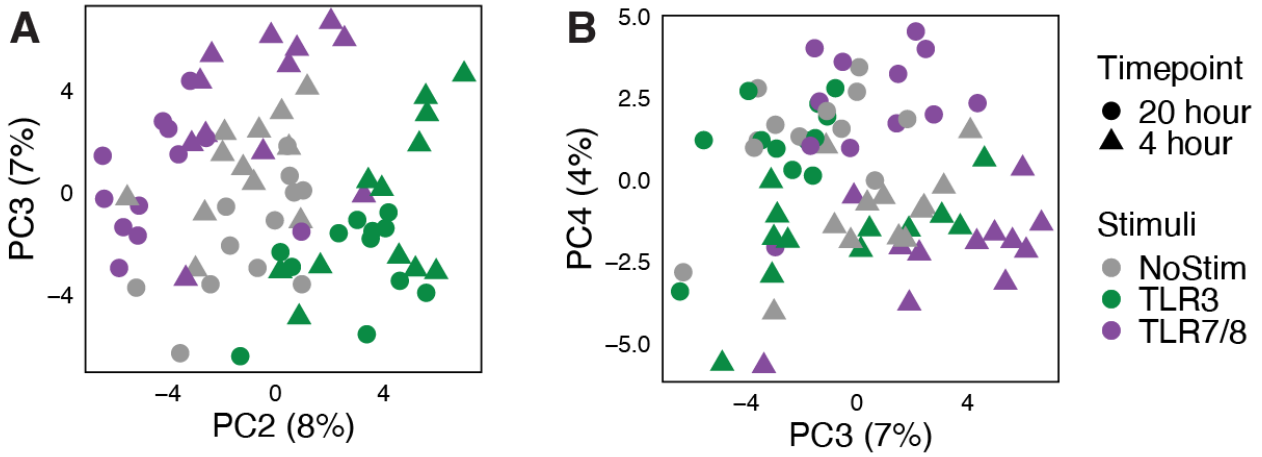


**Figure S5. PCA analysis of Olink protein expression results. (A & B)** Principal component analysis (PCA) of normalized protein expression (NPX) from unstimulated and TLR3- or TLR7/8-stimulated samples. PC3 and PC4 separate samples by time point.

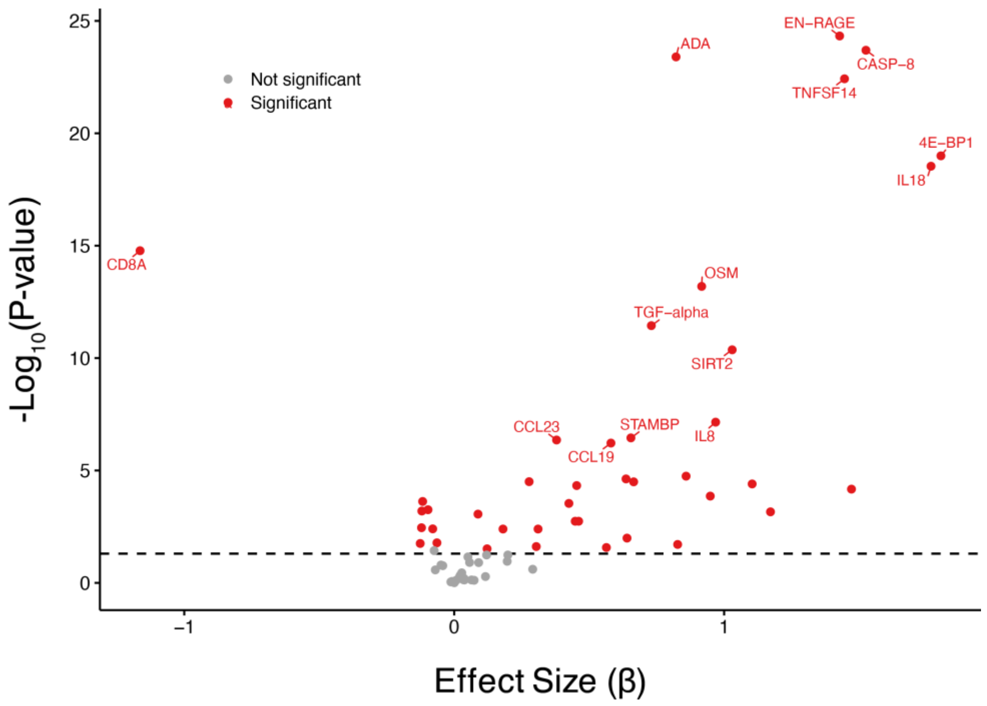


**Figure S6. Differential expression analysis results for protein expression over time.** Forty-one cytokines were differentially expressed at 20h vs 4h (FDR < 0.05), the majority of which (33 cytokines) showed increased expression with time. X axis: effect size, Y axis: log10p-value. The horizontal dashed line represents FDR = 0.05. Red dots show proteins that were significantly differentially expressed at 20 h vs 4 h (FDR < 0.05).

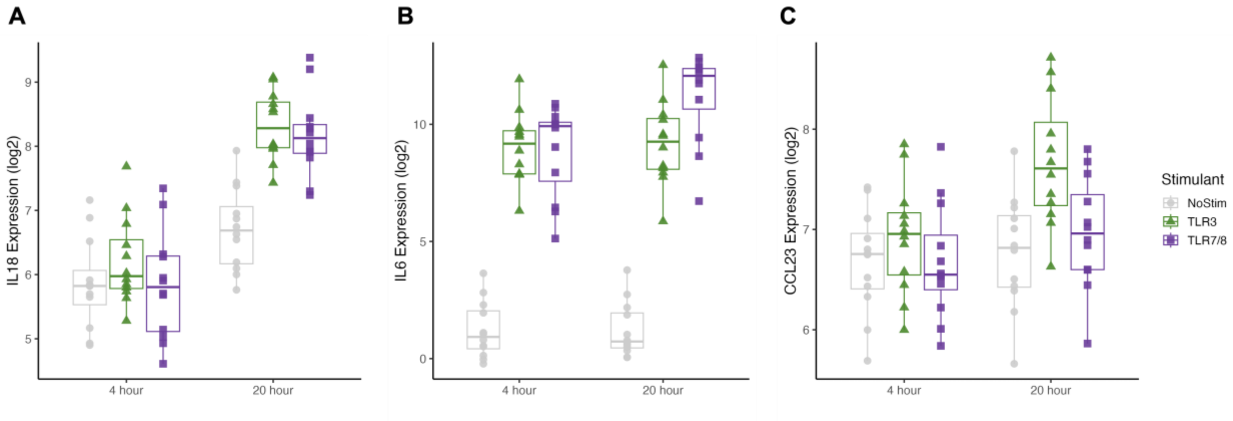


**Figure S7. Examples of cytokines that show stimulation-dependent time effects. (A)** IL18 shows stimulation-dependent time effects under both stimulation conditions. **(B)** IL6 exhibits stimulation-dependent time effects under TLR7/8 stimulation only. **(C)** CCL23 shows stimulation-dependent time effects under TLR3 stimulation only. X axis: time point, Y axis: cytokine expression ($log_2$).
